# Active degradation of a regulator controls coordination of downstream genes

**DOI:** 10.1101/272120

**Authors:** Nicholas A. Rossi, Thierry Mora, Aleksandra M. Walczak, Mary J. Dunlop

## Abstract

Several key transcription factors have unusually short half-lives compared to other cellular proteins. Here, we explore the utility of active degradation in shaping how a master regulator coordinates its downstream targets. We focus our studies on the multi-antibiotic resistance activator MarA, which controls a variety of stress response genes in *Escherichia coli*. We modify its half-life either by knocking down the protease that targets it via CRISPRi or by engineering MarA to protect it from degradation. Our experimental, analytical, and computational results indicate that active degradation can impact both the rate of coordination and the maximum coordination that downstream genes can achieve. Trade-offs between these properties show that perfect information fidelity and instantaneous coordination cannot coexist.

## Introduction

Active degradation is a rare feature in bacteria, affecting only 2-7% of total cellular proteins in *Escherichia coli* (Nath & Koch 1970; Maurizi 1992). Active degradation is accomplished by ATP-dependent proteases such as ClpXP, HflB, and Lon (Gottesman 2003). These proteases play a role in protein quality control by degrading misfolded proteins (Gur & Sauer 2008; Bezawork-Geleta et al. 2015; Laskowska et al. 1996), but they can also degrade functional proteins. Active degradation of functional proteins can serve as a type of post-translational regulation to modify protein concentration (Wickner et al. 1999). This can be important in time-sensitive processes when removal through dilution is insufficient. Metabolic processes that incorporate active degradation include DNA damage repair, gene expression during stationary-phase, cell division, filamentation, and accelerated removal of certain regulatory proteins (Fu et al. 1997; Flynn et al. 2003). Although active degradation can play a regulatory role, this comes with an energy cost due to the ATP necessary to activate the proteases, and a metabolic investment required to replace degraded proteins (Kafri et al. 2016). Active degradation occurs in many other microorganisms as well, such as with the ComK regulator of sporulation in *Bacillus subtilis* (Mugler et al. 2016) and in many regulatory proteins in humans including the p53 tumor suppressor (Schrader et al. 2009; Giaccia & Kastan 1998) and IκBα, which dictates NF-κB activity and cellular stress response (Mathes et al. 2008; Gilmore 2006). The refractory period of NF-κB pulses is determined by this degradation rate, which dictates the limit of information transmission in the system (Tudelska et al. 2017). In eukaryotes, the percentage of proteins that are primarily removed through active degradation ranges from ~15% in yeast (Christiano et al. 2014) to ~50% in humans (Eden et al. 2011). Therefore, despite active degradation’s role in regulation across domains of life, it is still comparatively rare in bacteria due to their rapid growth rate, which produces stronger dilution effects. The relative rarity coupled with the cost implicit to active degradation makes the presence of any short half-life protein in bacteria conspicuous. What potential utility could necessitate the use of this rare design feature?

To answer this, we focused on the multi-antibiotic resistance activator (MarA) in *E. coli* as a case study. MarA has a short half-life even by the standards of an actively degraded protein, with an estimated half-life of 1-3 minutes due to Lon protease activity (Griffith, Shah & E Wolf 2004a; Das et al. 2016). Moreover, MarA regulates over 60 downstream targets involved in a variety of antibiotic resistance roles, from genes encoding efflux pumps to small RNAs (Barbosa & Levy 2000; Martin et al. 1999). The majority of the downstream genes encode for stable proteins, therefore MarA has a much shorter half-life than most of its downstream targets (Barbosa & Levy 2000). Recent studies have revealed that *marA* is expressed stochastically, creating phenotypic diversity within isogenic populations, and this noisy expression is linked to transient antibiotic resistance (Meouche et al. 2016). Therefore, we asked what the potential advantages of active degradation of MarA are within the context of its stochastic dynamics and their propagation to downstream targets. In other words, we asked how MarA coordinates disparate genes necessary to create an antibiotic resistance phenotype.

Phenotypic diversity arises in isogenic populations due to stochastic expression of genes. The rate at which this diversity develops in growing populations depends on the time scale of the stochastic dynamics of the genes being expressed, including their rate of degradation (Garcia-Bernardo & Dunlop 2013; Mugler et al. 2016). Starting with a single cell at a single time point, there is no diversity. As cells grow and divide, diversity develops within the population. We consider diversity to be measurable as the distribution of distinct protein concentrations present in a population. The half-life of the protein impacts the rate of diversity generation and the maximum diversity achieved (Fig. 1A). For a regulatory protein, phenotypic diversity can propagate to its downstream targets (Fig. 1B). The rate at which diversity appears in downstream genes is impacted by how the upstream regulator changes with time. Active degradation affects the variability in the activator concentration—how quickly the concentration changes in one cell over time, as well as the diversity in concentrations between cells. This produces a spectrum of phenotypes where individual cells exhibit a range from low to high expression for all downstream targets (Fig. 1C). This coordinated diversity in expression of genes that share a master regulator is different from the uncoordinated diversity in two uncoupled genes (Fig. 1D). We used mutual information to quantify increasing coordination of downstream genes. Mutual information describes how well knowing the concentration of one of these proteins allows us to predict the concentration of the other (Tkacik & Walczak 2011). It measures their co-expression properties even if the correlation between them is nonlinear, an important feature because transcriptional dose response curves typically saturate, following the shape of a Hill function (Rossi & Dunlop 2017). Supplementary movie 1 shows an animation of how stochastic expression of an activator produces coordinated diversity within downstream targets.

**Figure 1.**
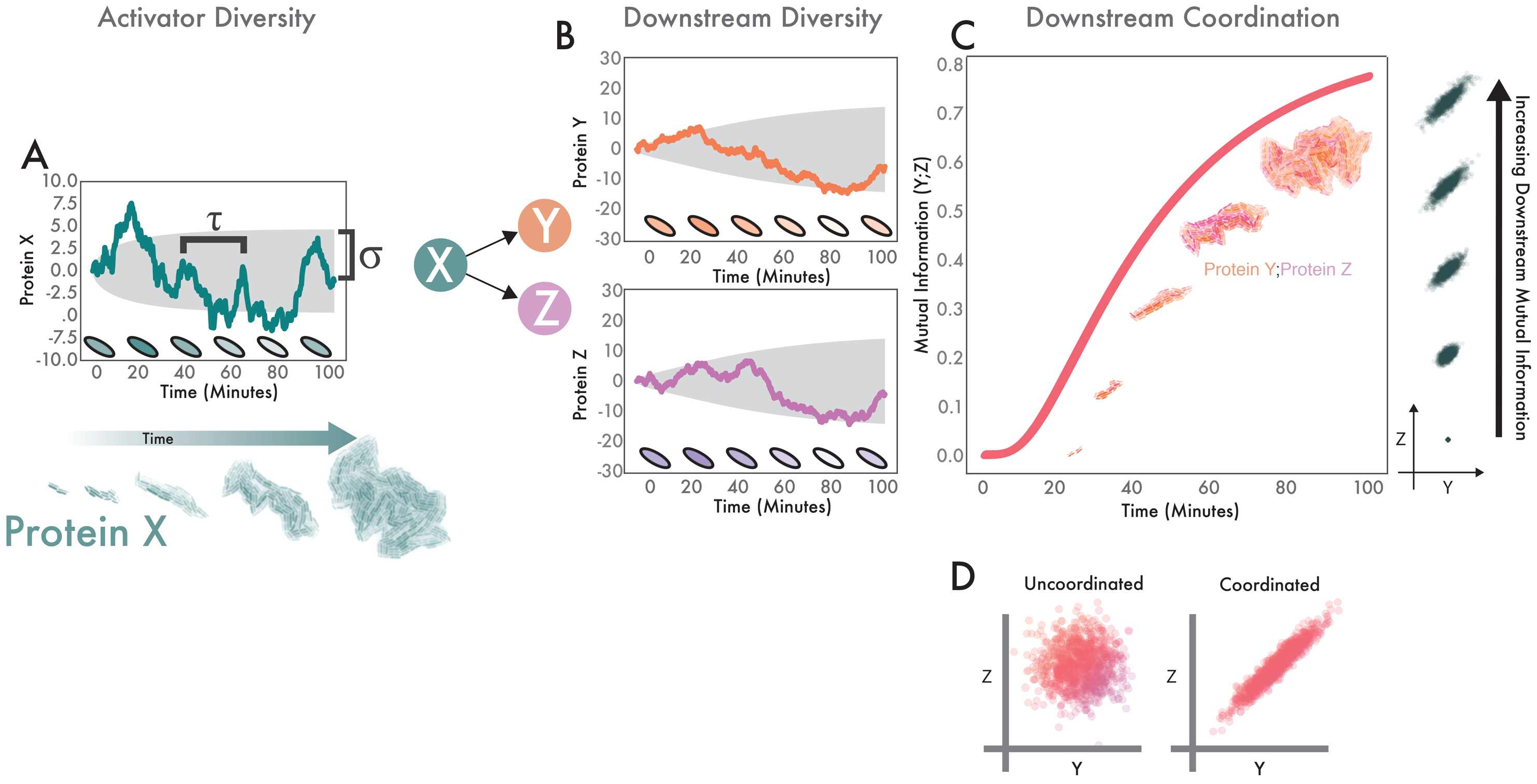
Statistics of stochastic protein expression in individual cells determine coordinated diversity in populations. **(A)** Variability in the master regulator (X) expands over time. Teal trace corresponds to a simulation of a single stochastic protein trajectory in one cell; shaded cells illustrate how we expect this to appear *in vivo*, τ governs how quickly the trace changes in time. Shaded gray region (σ) shows analytical solutions of how the projected diversity evolves over time (one standard deviation around the mean). Below, experimental data from growing microcolony shows how individual heterogeneity becomes population-level diversity. **(B)** Activator diversity propagates to downstream genes. Colored traces and gray shaded regions are the same as (A), but for Y and Z instead of X. **(C)** X produces growing coordinated diversity in Y and Z. Pink trace shows how coordinated diversity increases over time. Gray point clouds show how increasing information between two downstream targets appears visually. Experimental data from growing microcolony shows how heterogeneity between individual cells produces population-level variability. **(D)** Comparison between uncoordinated and coordinated downstream genes. Left scatter plot indicates two genes not governed by a master regulator. They have high variability, but low coordination. Right plot shows equally high variability, but increased coordination due to a common regulator.

Coordination as a function of a shared master regulator is important for establishing synergistic effects between downstream targets. For instance, MarA activates expression of genes in the AcrAB-TolC efflux pump. All three pump genes are necessary to export antibiotics. Upregulating one gene without the others may incur needless waste. Conceptually, this is like locking both the front door and the back door to a home in order to protect it; locking just one door does not make the home safer and simply expends energy. Thus, coordinating expression of stress response genes is essential.

Here, we investigate the role of activator half-life in controlling how a stochastic regulator generates coordinated diversity in multiple downstream genes. We do this by controlling the half-life of MarA, either by modulating the concentration of Lon protease or by protecting MarA itself from proteolytic degradation. We then evaluate the impact of MarA’s half-life by computing both the maximum amount of coordinated diversity that MarA can achieve as well as the rate at which it achieves it.

## Results

We first developed a computational model of how noise in MarA propagates to downstream genes as a function of its half-life to generate predictions that we could test experimentally. In the model, an upstream regulator (X) activates two downstream proteins (Y and Z). We modeled noise in X using an Ornstein-Uhlenbeck process (Gillespie 1996; Rossi & Dunlop 2017; Dunlop et al. 2008). The model has two free parameters: τ, which is a scaled version of the half-life of the molecule (τ = λ/ln(2), where λ is the half-life) and σ, which specifies the noise in protein expression. Using this model, τ changes the dynamics of the process as well as the population distribution that such a process can produce (Fig. 2A). The stochastic simulations demonstrate that having a longer half-life causes the protein concentration to change more slowly. Additionally, the longer half-life allows the trajectory of protein concentration to wander farther away from the mean, increasing the standard deviation (Fig. 2A). In this case, the standard deviation scales as 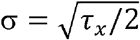 (Gillespie 1996). Analytical solutions to these functions show that as the half-life of a molecule increases, it takes longer to reach the maximum variability (Fig. 2B).

**Figure 2.**
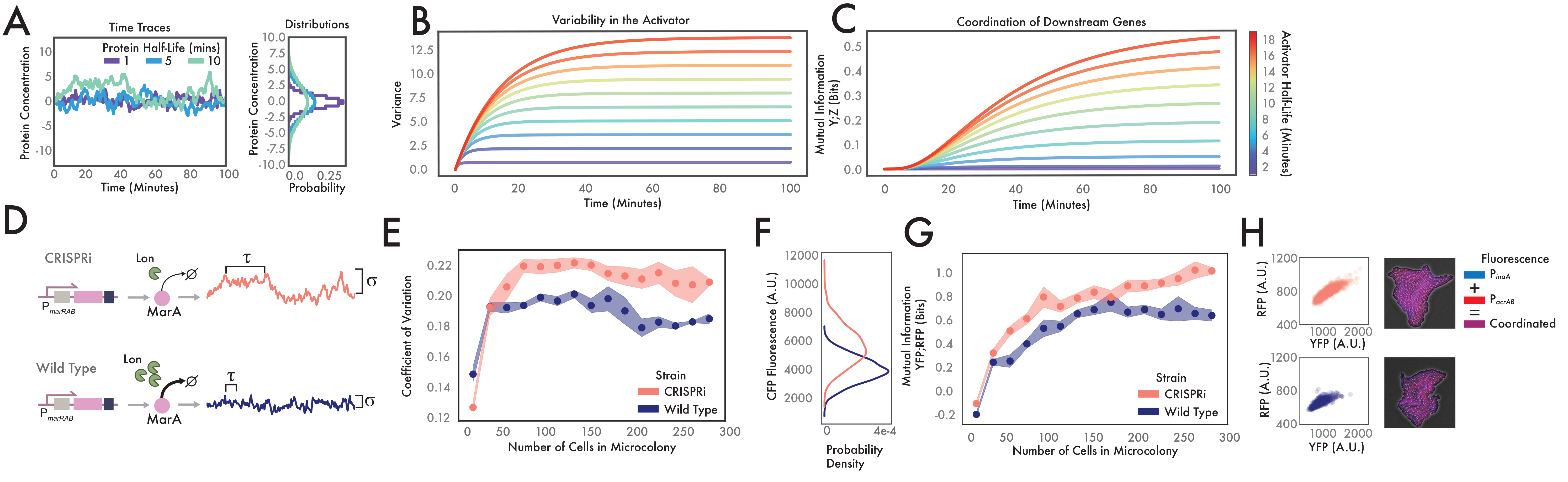
CRISPRi knockdown of Lon protease to increase half-life. **(A)** Simulation of how the half-life of regulator X affects correlation time and variability. Sample trajectories with varying degradation rates produce different probability distributions (right). **(B)** Analytical solutions for the variability of a protein as a function of differing half-lives. Individual traces demonstrate increasing variance as a function of time and the half-life. **(C)** Analytical solutions for the coordination of two downstream genes as a function of differing activator half-life. Individual traces demonstrate increasing coordination in two downstream genes as a function of time and the half-life of the activator. **(D)** Experimental schematic for CRISPRi knockdown of Lon protease. Upper row shows how time-constant τ and standard deviation σ are modified via decreased concentration of Lon protease due to CRISPRi knockdown. Bottom row indicates wild type system where Lon actively degrades MarA, producing time-series changes in protein concentration. **(E)** Experimental results showing differences in activator variability between wild type and CRISPRi systems. Dots show mean coefficient of variation over n=5 growing microcolonies. Shaded regions show standard error (Fig. S2). **(F)** Histograms showing distributions of *P*_*marA*_ at final time point of (E). **(G)** Downstream coordination expands differently in wild type and CRISPRi strains. Mutual information between YFP and RFP (P_*inaA*_ and *P*_*acrAB*_) is calculated across replicates in n=5 growing microcolonies. Mean and standard error are represented as in (E). **(H)** Maximum coordinated diversity is increased in CRISPRi system over wild type. Bivariate scatter plots show P_*inaA*_ and P_*acrAB*_ expression levels at the final time point across n=5 microcolonies. Snapshots give examples of coordination in a single microcolony. Blue is P_*inaA*_, red is P_*acrAB*_, downstream coordination appears as magenta. Note that lack of coordination appears as red or blue cells.

Variability in a regulatory molecule propagates to the downstream genes it controls. To quantify the correlation between the concentrations of two downstream proteins (Y and Z) regulated by X, we measured the mutual information between their concentrations as a function of time. First, we observed an apparent advantage to longer half-life activators, as increasing half-life increases the mutual information between two downstream targets (Fig. 2C). Large variability in the activator concentration allows for coordinated diversity in the downstream genes, and a longer half-life molecule produces greater information between the two downstream genes. The decreased mutual information of short half-life activators is due to filtering of high-frequency changes. Downstream genes function as low-pass filters in which signals that change quickly are averaged out (Hooshangi et al. 2005). Moreover, because increasing the half-life increases the standard deviation of protein levels, the variability in downstream genes increases as well, allowing for a greater range of coordinated activation. These results show that longer half-life proteins may be better at generating coordinated diversity in downstream targets. However, this conclusion is dependent on allowing the standard deviation to scale freely with the half-life of the regulatory molecule.

To validate these analytic and computational predictions, we designed a genetic system to modify the concentration of Lon protease. Changing the concentration of Lon should change the statistics of MarA and the downstream genes it regulates. We used CRISPRi to knock down Lon expression, allowing us to control the level of MarA. We first transformed *E. coli* with dCas9 and a sgRNA targeting *lon* (Qi et al. 2013). Lon protease knockouts have been shown to increase the mean concentration MarA in *E. coli*, but affect many other genes in addition to *marA* (Martin et al. 2008; Griffith, Shah & E Wolf 2004a). In order to avoid off-target pleotropic effects from eliminating Lon protease such as cell filamentation (Schoemaker et al. 1984), we designed the sgRNA to increase MarA levels, but to produce no qualitative morphological changes to the cell (Fig. S1). Decreasing Lon protease is expected to affect the correlation time τ of MarA, as well as the standard deviation o of its time trace (Fig. 2D).

We co-transformed cells with the CRISPRi construct with a plasmid containing transcriptional reporters for MarA and its downstream genes. To report MarA levels, the plasmid contains a modified *marRAB* promoter controlling cyan fluorescent protein (*cfp*). Our analytical results predict that extending the half-life of an activator should produce increased variability in growing microcolonies. By measuring the CFP levels of individual cells in the CRISPRi knockdown system and a wild type strain we found that the half-life of the molecule determines how noise expands in growing microcolonies (Fig. 2E). The analytical results predict that the rate of diversity generation should not be altered in growing microcolonies, but that the maximum diversity is (as quantified by coefficient of variation). Consistent with this, we found that the slope of the curves was similar, but the magnitude changes. The distribution of CFP within microcolonies at the final time step of the experiments highlights the differences in maximum diversity (Fig. 2F). As in the theoretical predictions, the mean and standard deviation of MarA in the CRISPRi knockdown system, which has a longer half-life, increase compared to the wild type system.

In addition to the *marA* reporter, we also measured expression of two downstream genes: *inaA*, which is a pH-inducible gene involved in stress response (Rosner & Slonczewski 1994) and *acrAB*, which is a component of the *acrAB-tolC* antibiotic efflux pump (Du et al. 2014; Okusu et al. 1996). We selected these two genes because of their disparate functionality, yet similar dose response curves (Rossi & Dunlop 2017). We used P_*inaA*_ to control expression of yellow fluorescent protein (*yfp*) and P_*acrAB*_ to control red fluorescent protein (*rfp*). By simultaneously measuring the dynamics of all three fluorophores (P_*marA*_-*cfp*, P_*inaA*_-*yfp*, P_*acrAB*_-*rfp*) via time-lapse microscopy, we were able to quantify variability in the *marA* input and two outputs in both the CRISPRi and wild type systems.

We used these fluorescence data to calculate the coordinated diversity in the microcolony by measuring mutual information over time. This metric captures the value of a large spectrum of correlations between two downstream genes without over-valuing high correlations between a few cells, such as would occur during the beginning growth stages of a microcolony (Tkacik & Walczak 2011). Mirroring the analytical results, we see that there is an advantage to increasing the half-life of MarA, where coordinated phenotypic diversity is higher between the two downstream genes in the CRISPRi strain (Fig. 2G).

To emphasize the differences, we compared single-cell data from the final time point of the movies. As expected, the mean concentration of both YFP and RFP increases in the CRISPRi system relative to wild type due to decreased MarA degradation (Fig. 2H). Importantly, sample snapshots of the two conditions show greater correlation between the fluorophores (higher proportion of magenta cells) in the longer half-life CRISPRi system. This illustrates that in addition to increasing mean expression of the downstream genes, the correlation between downstream gene promoter activities increases with the half-life of the activator.

These results show clear advantages for longer half-life activators (equivalent to conditions without active degradation), but they do so without constraints – the variance of the activator is allowed to increase without a limit. This may not be the case in natural systems which have a target concentration range for the activator molecule. In order to investigate this, we modified our mathematical model to create a system where the standard deviation does not increase as a function of the activator half-life. We did this by requiring the noise to be inversely related to the correlation time 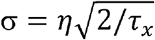. This represents a system where the total variability is constrained, but where the rate at which it grows is a free parameter. Three example trajectories from stochastic simulations with varying degradation rates but normalized standard deviations show different time series behaviors that produce identical steady state distributions (Fig. 3A). Under this constraint, we see that the rate at which the variance of a population develops is a function of activator half-life (Fig. 3B). Moreover, when the activator variance is constrained, a short halflife activator increases mutual information faster, but plateaus sooner and at lower values than a longer half-life activator (Fig. 3C). Thus, faster-degrading activators are capable of generating coordinated diversity faster, but the maximum correlation that they can produce between two downstream genes is limited. Here is the first evidence of a potential trade-off as we increase activator half-life by inhibiting active degradation: maximum coordination between two downstream targets increases but the rate at which coordination is generated decreases.

**Figure 3.**
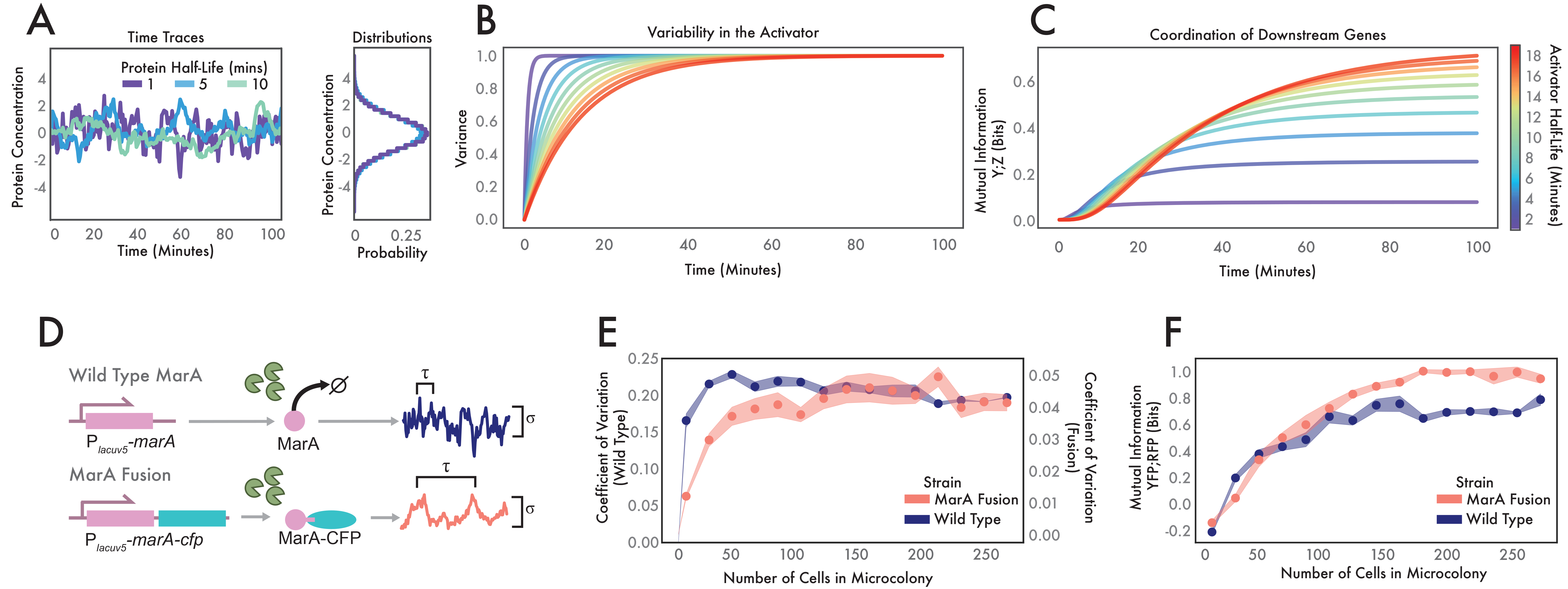
MarA-CFP fusion shows different downstream statistics than wild type MarA, even at similar expression levels. **(A)** Simulation of how modulating the half-life of regulator X affects correlation time. Sample trajectories with varying half-lives but constrained population variance produce the same distributions (right). **(B)** Analytical solutions for the variability of a protein as a function of time and half-life. **(C)** Analytical solutions for the coordination of two downstream genes as a function of differing activator half-lives. **(D)** Experimental schematic comparing wild type and MarA fusion systems. Top row indicates wild type system. Bottom row indicates protection of MarA from active degradation by Lon via a translational fusion with CFP. This modification changes the correlation time τ compared to wild type. IPTG induction of the LacUV5 promoter allows us to adjust σ independently of τ. **(E)** Experimental results showing growing variability of wild type and MarA fusion strains. **(F)** Experimental results of growing coordinated diversity of wild type and MarA fusion. For (E) and (F) dots show mean coefficient of variation over n=5 growing microcolonies. Shaded regions show standard error.

In order to verify these theoretical results experimentally we developed a system in which we could control the mean activator concentration independently of the half-life. To this end, we employed a modified version of MarA that was resistant to degradation by Lon and placed it under the control of an inducible promoter. In this design, *marA* is translationally fused to *cfp* thereby protecting the C terminus from recognition and degradation by Lon protease (Meouche et al. 2016; Smith et al. 1999). Expressing the MarA fusion from the P_*lacUV5*_ promoter allowed us to tune its expression level using IPTG. Thus, we were able to tune the steady state concentration of MarA and the noise that scales with it. We transformed the MarA fusion plasmid into *E. coli* Δ*marRAB* along with a two-color version of the downstream gene reporter plasmid (P_*lacUV5*_-*marA-cfp* with P_*inaA*_-*yfp*, P_*acrAB*_-*rfp*). By monitoring the real-time MarA level directly using CFP, we were able to quantify how much MarA was present in each cell. We compared this strain with altered MarA degradation to a system with wild type MarA under the control of the P_*lacUV5*_ promoter as well as the three-color reporter described previously (P_*lacUV5*_-*marA* with P_*marA*_-*cfp*, P_*inaA*_-*yfp*, P_*acrAB*_-*rfp*).

The two systems produce protein concentrations with different statistics over time (Fig. 3D). The MarA fusion has a longer correlation time due to protection from Lon protease, however unlike in the CRISPRi system, the expression level can be tuned via IPTG induction. In experiments, we observed that wild type MarA achieved its maximum diversity faster than the MarA fusion (Fig. 3E). This is in contrast to the CRISPRi knockdown system in which the rates did not change substantively as a function of half-life. This is an essential difference because the rate of activator diversity generation determines the rate of coordination in downstream targets. Supplementary movies 2 and 3 show how the growing microcolonies exhibit differences in how diversity is generated. Wild type MarA produces diversity quickly while the microcolony expressing the MarA fusion takes longer. Therefore, we see that both maximum diversity and the rate of diversity generation can be altered by modulating the half-life of the molecule.

As with the CRISPRi system, the changes in activator dynamics go on to alter how downstream genes are regulated, where increasing MarA’s half-life increases the maximum coordinated diversity between downstream genes (Fig. 3F). However, unlike in Fig. 2G, the rate at which the wild type and MarA fusion generate coordination between downstream genes is different. We observed that the wild type strain approached its maximum degree of coordination between the two genes faster than the MarA fusion. This is a reflection of activator diversity accumulating faster when the system produces wild type MarA. Also, in contrast to the CRISPRi knockdown example, here both strains express the same steady state levels for each of the downstream genes. This shows that even if the mean expression is equivalent, the coordination between genes can be different (Fig. S3).

So far, we have compared the maximum coordinated diversity that a system can produce and the rate at which this diversity grows. The information rate is an important metric to consider because it explicitly outlines the role that a stochastic activator plays in managing the progressive development of phenotypic diversity among multiple genes. In order to conceptualize the difference between the information rate and total mutual information, consider Fig. 4A showing two increasingly resolved maple leaves. We see that in the two rows, we must consider both the rate at which the image gains clarity as well as the maximum clarity of the image. Genetic networks may optimize for increased rate at the expense of low mutual information in time-sensitive conditions. Alternatively, a decreased information rate may be acceptable if time constraints are not an issue and total information is more important. To quantify this trade-off, we used the methods described in (Tostevin & Wolde 2009) to calculate the information rate for our model and compared that to the maximum information between two downstream genes. First, at extremely short half-lives both the information rate and the maximum information are very low. This is because near the uncorrelated white noise limit, the activator signal X is filtered by the downstream genes Y and Z. However, as we increase the activator half-life we see a marked increase in both information rate and mutual information. This is a function of a rapidly changing activator passing information to downstream targets, which allows them to coordinate. As the activator half-life increases further, the maximum information continues to increase monotonically while the information rate decreases. A slowly changing signal in X is passed faithfully to downstream targets but the rate of information is slow. Note however that information transmission rate is not simply the derivative of mutual information over time, as it is computed from the complete power spectra of the two signals and includes information passed with time delays (Tostevin & Wolde 2009) (Methods).

**Figure 4.**
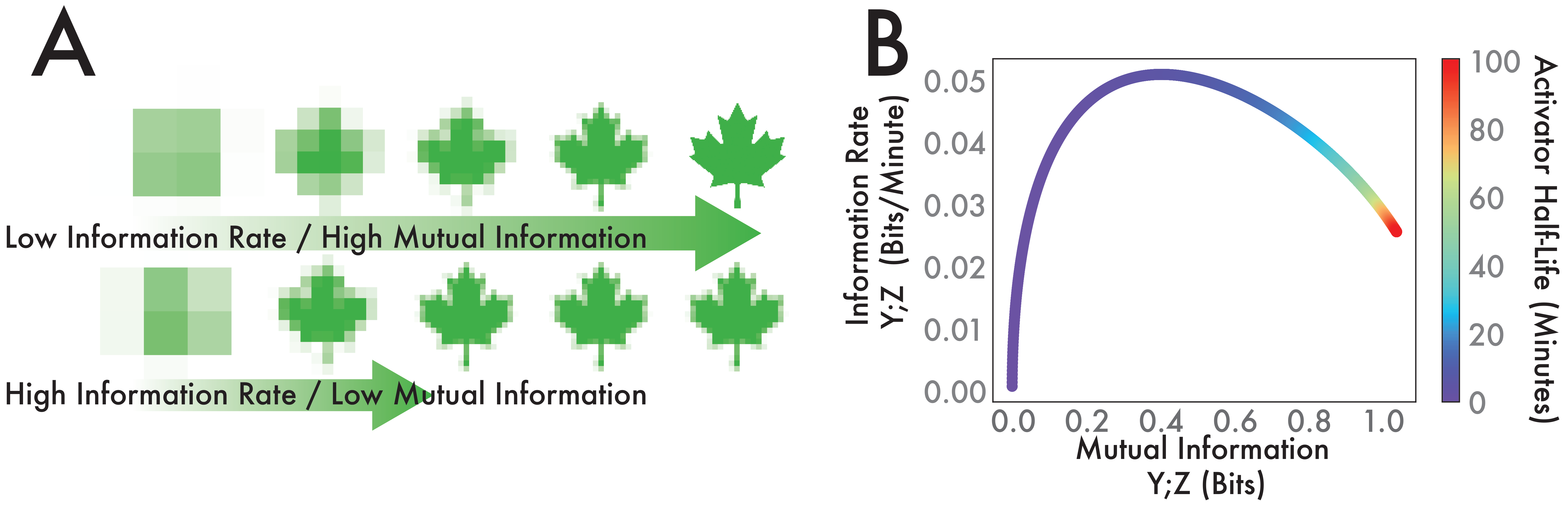
Trade-offs in mutual information versus information rate as a function of activator half-life. **(A)** Illustration of the differences between information rate and maximum information. Growing resolution of maple leaf indicates increased information. Top row has higher maximum information, but it is accrued at a lower rate as compared to the bottom row. **(B)** Mutual information and information rate between two downstream genes as a function of activator half-life.

We have demonstrated how active degradation of a master regulator can change the way a growing microcolony generates coordinated phenotypic diversity. Importantly, we have highlighted a role for active degradation in increasing the rate of information transmission. However, this increased information rate comes at the cost of decreased maximum mutual information between downstream targets. Together, these opposite design demands represent a trade-off that genetic networks can optimize based on individualized requirements.

## Discussion

In *E. coli*, several key regulators are subject to active degradation. Here, we demonstrate a role for active degradation using MarA as a case study. We show that the half-life of a regulatory molecule affects not only its time-series behavior and population statistics, but also its ability to coordinate downstream genes. This downstream coordination develops over time, with both a rate and maximum value.

We focus on how individual founders of new bacterial populations can quickly establish a spectrum of coordinated phenotypes. We show that generating variability in downstream targets is limited by variability in the activators that control them. Also, signals from activators that change too quickly can be filtered, narrowing the spectrum of coordination in downstream targets. Our results indicate that there is a trade-off between information rate and maximum information in coordinating downstream targets. While this will depend on a number of system specific parameters, this trade-off shows that perfect information fidelity and instantaneous coordination cannot coexist. This is analogous to instructing two people to do a task in tandem-the faster you speak, the faster they can be instructed to coordinate and execute the objective, but at some point information is lost and mistakes will be made.

Although we focused on how individual genes and proteins generate phenotypic diversity, extrinsic factors like growth rate have the potential to affect this as well. For instance, varying the cell division time can affect how variability in MarA forms (Fig. S4). In effect, the analytical solutions to the variance functions represent the upper limit on how diversity develops, and decreasing the growth rate scales this curve.

While our experiments and models focused on downstream genes where the typical concentrations of MarA fall within their linear regime of activation, it should be noted that this is not true for all downstream targets of MarA. Previous research has highlighted that each downstream target possesses a unique nonlinear dose response curve as a function of MarA activation (Rossi & Dunlop 2017; Martin et al. 2008). It may be possible for growing cells to exploit the inherent nonlinearity of each downstream gene’s activation curve to better achieve the stoichiometric balance required by downstream targets with wide ranging functions. That is to say, while multiple downstream targets may be necessary to mount an effective barrier to stressors such as antibiotic challenge, the optimal activation curves of each may be unique. This flexibility of downstream activation allows for this balance to occur despite signals coming from a single shared activator. Moreover, we have focused on the role of MarA as a stochastic activator, however the concentration of MarA is also dependent on the environment (Hao et al. 2014). The utility of active degradation may play a role in signal transmission as well, decreasing the response time (Alon 2007; Griffith, Shah & E Wolf 2004b).

Using a combination of single-cell time lapse movies with multiple fluorescent reporters, analytical models, and computational simulations we have demonstrated a role that protein halflife plays in generating single-cell variability and the ability of an activator to produce coordinated diversity in downstream genes. Our results show that both maximum coordinated diversity as well as the rate of coordination between multiple targets is dependent on activator half-life. This suggests a possible design advantage for active degradation and sheds light on how cells grow from individuals to diverse populations, while coordinating expression of related processes.

## Methods

### Strains and plasmids

We used two strains for the experiments: wild type *E. coli* MG1655 and *E. coli* MG1655 Δ*marRAB* from (Meouche et al. 2016).

In order to construct the three color reporter plasmid, we modified pNS2-σVL from (Dunlop et al. 2008) by placing P_*marA*_, P_*inaA*_, and P_*acrAB*_ upstream of cyan fluorescent protein (CFP), venus yellow fluorescent protein (YFP), and mCherry red fluorescent protein (RFP), respectively. All other plasmids were derived from the BioBrick library described in (Lee et al. 2011).

For the CRISPRi knockdown of Lon protease, we transformed the plasmid system from (Qi et al. 2013), using one plasmid containing the constitutively expressed Lon targeting sgRNA and another plasmid containing dCas9 under the control of a tetracycline inducible promoter.

For the MarA fusion experiments, we cotransformed the IPTG inducible MarA-CFP translational fusion plasmid from (Meouche et al. 2016) with a two-color reporter plasmid bearing P_*inaA*_ and P_*acrAB*_ controlling YFP and RFP into *E. coli* Δ*marRAB*.

Additional details on plasmid construction are available in Supplementary Information.

### Time-lapse fluorescence microscopy

Cultures were inoculated from single colonies and grown overnight at 37°C with 200 rpm shaking in Luria-Broth (LB) medium. All strains were grown in 30 μg/ml kanamycin. In addition, strains containing the CRISPRi knockdown system were grown in the presence of 100 μg/ml carbenicillin and 30 μg/ml chloramphenicol. Strains containing the wild type or MarA fusion inducible plasmids were grown with 100 μg/ml carbenicillin in addition to kanamycin.

Overnight cultures were diluted 1:100 in selective LB medium. For the wild type or inducible MarA experiments, we added 0 μM IPTG for the MarA-CFP fusion and 10 μM IPTG for wild type MarA and grew cultures for four hours. The differences in IPTG induction allowed for similar mean concentrations with different degradation kinetics (Fig. S3). For the other experiments containing CRISPRi knockdown plasmids, cultures were refreshed for 4 hours without addition of inducer. For microscopy movies, we placed cells on 1.5% MGC low melting point agarose pads (Dunlop et al. 2008). We used a Nikon Instruments Ti-E microscope to image cells at 100× magnification. Time-lapse movies were taken at a temporal resolution of every three minutes for 10 hours. We used Super Segger cell tracking software to extract fluorescence data from individual cells (Stylianidou et al. 2016). When computing the coefficient of variation and the mutual information of growing microcolonies of a given size, data from five replicates was combined and binned according to the number of cells in the microcolony (Fig. S2).

### Stochastic simulations

We modeled the activator X and the downstream products Y and Z by:

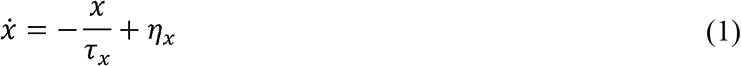

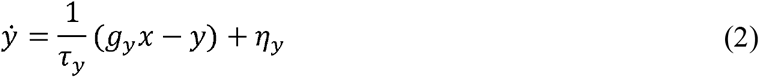

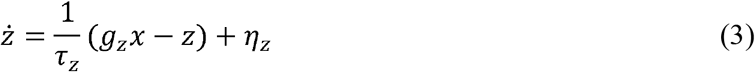

where X represents the activator concentration and Y and Z are the concentrations of two downstream proteins. This mirrors P_*marRAB*_, P_*inaA*_, and P_*acrAB*_ in our experimental system. τ is the correlation time of each of the downstream proteins and is proportional to the half-life (λ) of the molecule: τ = λ/ln(2). g is the gain of each downstream promoter. *σ* is the noise scaling term, which is set equal to one in the case where the variance is not normalized by the correlation time or is allowed to inversely scale with the correlation time: 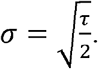 η is a zero mean Gaussian distributed white noise random variable.

### Variance

To compute the analytical solutions to the variance over time we applied the methods from (28) to Eqn. 1:

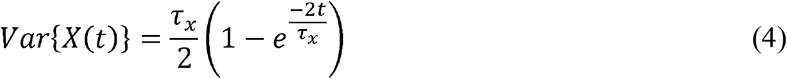

To compute the variance for systems that converge to the same final value, we modified Eqn. 1 to include a normalization term:

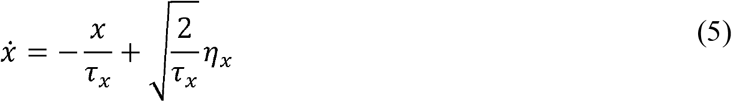

Computing the variance over time from this function yields:

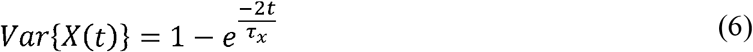

Derivations for variance calculations of downstream genes can be found in Supplementary Information.

### Mutual Information

Mutual information is calculated from the analytical solution to the correlation function between Y and Z:

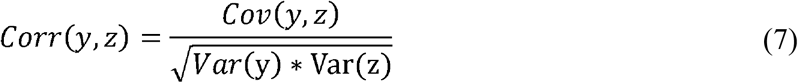

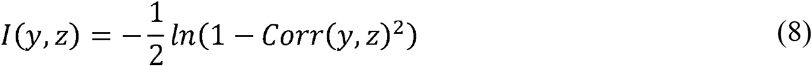

The maximum mutual information between Y and Z is computed analytically by calculating the correlation using Eqns. 2 and 3.

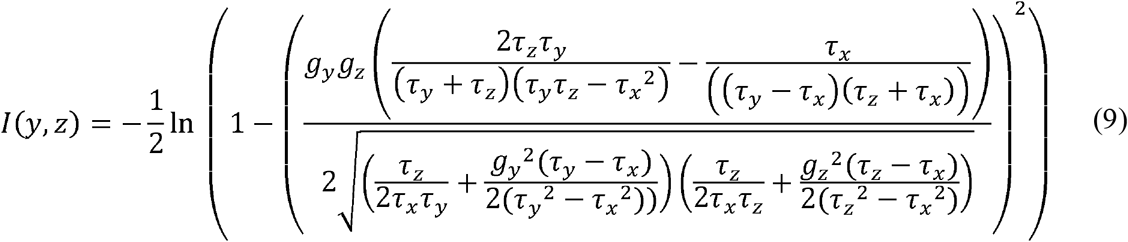

The analytical calculations for mutual information over time in growing microcolonies can be found in Supplementary Information.

Mutual information was calculated from experimental fluorescence data using the YFP and RFP fluorescence data and k-nearest neighbor distances (k=3) (Kraskov et al. 2004).

### Information rate

The information rate between downstream genes is computed using Eqns. 2 and 3 and following the methods outlined in (Tostevin & Wolde 2009). S_yz_ is the cross power spectral density between Eqns. 2 and 3, S_yy_ is the power spectral density for Eqn. 2, and S_zz_ is the power spectral density for Eqn. 3:

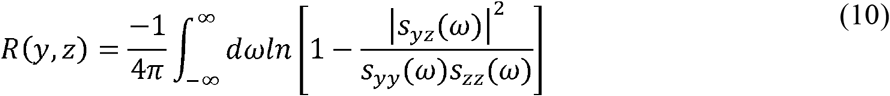

Computing the above power spectra and performing the integration yields the equation:

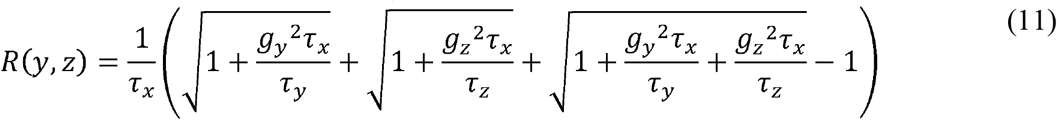

**Figure S1. Morphology of wild type, Δ*lon*, and CRISPRi knockdown strains**

Phase contrast images of microcolonies after 200 minutes of growth. Slower growth rate and increased filamentation are evident in the Δ*lon* strain.

**Figure S2. Generating coefficient of variation statistics from growing microcolonies.**

Microcolonies were allowed to grow for 300 minutes with data taken every 3 minutes. Variance was computed for each microcolony at each time point and then plotted against the number of cells in the microcolony (orange dots). These data were then binned (gray bars) and summary statistics such as the mean coefficient of variation across microcolonies (green dots) were generated for each bin.

**Figure S3. Downstream gene expression for wild type and MarA fusion strains.**

YFP (P_*inaA*_) and RFP (P_*acrAB*_) fluorescence values from growing microcolony data plotted without the time dimension. Error bars show means and standard deviations for the two bivariate distributions. 10 μM IPTG induced P_lacuv5_-MarA elicits a very similar downstream response to 0 μM IPTG induced P_lacUV5_-MarA-CFP translational fusion (two distributions are statistically equivalent by the 2D Kolmogorov-Smirnov test, p>0.1).

**Figure S4. Impact of growth rate on generation of MarA variance.**

**(A)** Example simulation of a growing bacterial microcolony with a cell division time of 15 minutes. Individual trajectories show level of X.

**(B)** Analytical solution to the variance functions including growth rate terms. Plot also shows the average variance for 1000 stochastic simulations with cell division. The shaded regions represent the standard error over all simulations centered around the mean for each simulation set. The solid lines represent the analytical solutions, with the gray line representing the theoretical maximum. See Supplementary Information for growth rate functions.

**Supplementary Movie 1: Animation of activator noise propagating to downstream targets to produce coordinated diversity.**

**(A)** Example of stochastic activator (X) expression over time. Teal line shows example trajectory; shaded gray region outlines one standard deviation around the mean.

**(B)** Orange line shows example trajectory of downstream gene Y as it responds to signal from X. Orange dots show location of other example trajectories over time. Shaded gray region outlines analytical solution for standard deviation around the mean.

**(C)** Purple line shows example trajectory of downstream gene Z as it responds to signal from X. Purple dots show location of other example trajectories over time. Shaded gray region outlines analytical solution for standard deviation around the mean.

**(D)** Analytical solution to mutual information over time. Red dot shows current mutual information at time point in simulation.

**(E)** Visualization of mutual information between downstream genes. Each gray dot shows example trajectory value between Y and Z for a given time point of the simulation. Orange and purple dots from B and C are examples of the dots here.

**Supplementary Movie 2: Diversity in wild type MarA in a growing microcolony**

**(A)** Example of a growing bacterial microcolony expressing CFP under control of the P_*marA*_ promoter.

**(B)** Time traces of fluorescence for each cell in the microcolony.

**Supplementary Movie 3: Diversity in MarA fusion in a growing microcolony**

**(A)** Example of a growing bacterial microcolony with the MarA-CFP fusion strain.

**(B)** Time traces of fluorescence for each cell in the microcolony.

## Acknowledgments

We thank Nikit Patel and Imane El Meouche for their critical reading of the manuscript. This research was supported by the National Science Foundation (grant No. 1347635, MJD), the National Institutes of Health (grant No. 1R01AI102922, MJD), and a Chateaubriand Fellowship (NAR).

## Author Contributions

Conceptualization: NAR, TM, AMW, MJD. Investigation: NAR. Methodology: NAR, TM, AMW, MJD. Writing: NAR, MJD.

